# A Deep Learning method for classification of HNSCC and HPV patients using single-cell transcriptomics

**DOI:** 10.1101/2023.08.24.554735

**Authors:** Akanksha Jarwal, Anjali Dhall, Akanksha Arora, Sumeet Patiyal, Aman Srivastava, Gajendra P. S. Raghava

## Abstract

Head and Neck Squamous Cell Carcinoma (HNSCC) is the seventh most highly prevalent cancer type worldwide. Early detection of HNSCC is one of the important challenges in managing the treatment of the cancer patients. Existing techniques for detecting HNSCC are costly, expensive, and invasive in nature. In this study, we aimed to address this issue by developing classification models using machine learning and deep learning techniques, focusing on single-cell transcriptomics to distinguish between HNSCC and normal samples. Additionally, we built models to classify HNSCC samples into HPV-positive (HPV+) and HPV-negative (HPV-) categories. The models developed in this study have been trained on 80% of the GSE181919 dataset and validated on the remaining 20%. To develop an efficient model, we performed feature selection using mRMR method to shortlist a small number of genes from a plethora of genes. Artificial Neural Network based model trained on 100 genes outperformed the other classifiers with an AUROC of 0.91 for HNSCC classification for the validation set. The same algorithm achieved an AUROC of 0.83 for the classification of HPV+ and HPV-patients on the validation set. We also performed Gene Ontology (GO) enrichment analysis on the 100 shortlisted genes and found that most genes were involved in binding and catalytic activities. To facilitate the scientific community, a software package has been developed in Python which allows users to identify HNSCC in patients along with their HPV status. It is available at https://webs.iiitd.edu.in/raghava/hnscpred/.

**Key Points:**

- Application of single cell transcriptomics in cancer diagnosis
- Development of models for predicting HNSCC patients
- Classification of HPV+ and HPV-HNSCC patients
- Identification of gene biomarkers from single cell sequencing
- A standalone software package HNSCpred for predicting HNSCC patients

**Author’s Biography:**

1. Akanksha Jarwal is currently pursuing an M. Tech. in Computational Biology at the Department of Computational Biology, Indraprastha Institute of Information Technology, New Delhi, India.
2. Anjali Dhall is currently pursuing a Ph.D. in Computational Biology at the Department of Computational Biology, Indraprastha Institute of Information Technology, New Delhi, India.
3. Akanksha Arora is currently pursuing a Ph.D. in Computational Biology at the Department of Computational Biology, Indraprastha Institute of Information Technology, New Delhi, India.
4. Sumeet Patiyal is currently pursuing a Ph.D. in Computational Biology at the Department of Computational Biology, Indraprastha Institute of Information Technology, New Delhi, India.
5. Aman Srivastava is currently pursuing an M. Tech. in Computational Biology at the Department of Computational Biology, Indraprastha Institute of Information Technology, New Delhi, India.
6. Gajendra P. S. Raghava is currently working as a Professor and Head of the Department of Computational Biology, Indraprastha Institute of Information Technology, New Delhi, India.

## Introduction

Head and neck cancer, encompasses a variety of malignancies that affect the respiratory tract and upper digestive tract. Head and Neck Squamous Cell Carcinoma (HNSCC) is the most typical kind among the head and neck cancer [1]. In 2020, 562,328 people were diagnosed with head and neck cancer (HNC) worldwide, with a total count of 277,587 deaths due to the disease [2]. These carcinomas often develop in the salivary glands, larynx, oral cavity, throat, and sino-nasal tract epithelium. A number of head and neck malignancies are linked to the human papillomavirus (HPV) infection, notably HPV-16. However, some malignancies are also related to the other carcinogens like smoking, excessive alcohol, and other factors depending on the country or area. Hence, we can classify this cancer into two major categories - HPV-negative and HPV-positive. The median age of diagnosis for HPV associated HNSCC is about 66 years, whereas for HPV-associated oropharyngeal cancer the median age is ∼53 years [3]. The mechanisms of HPV+ and HPV-associated HNSCC are explained in Figure 1.

**Figure 1:**
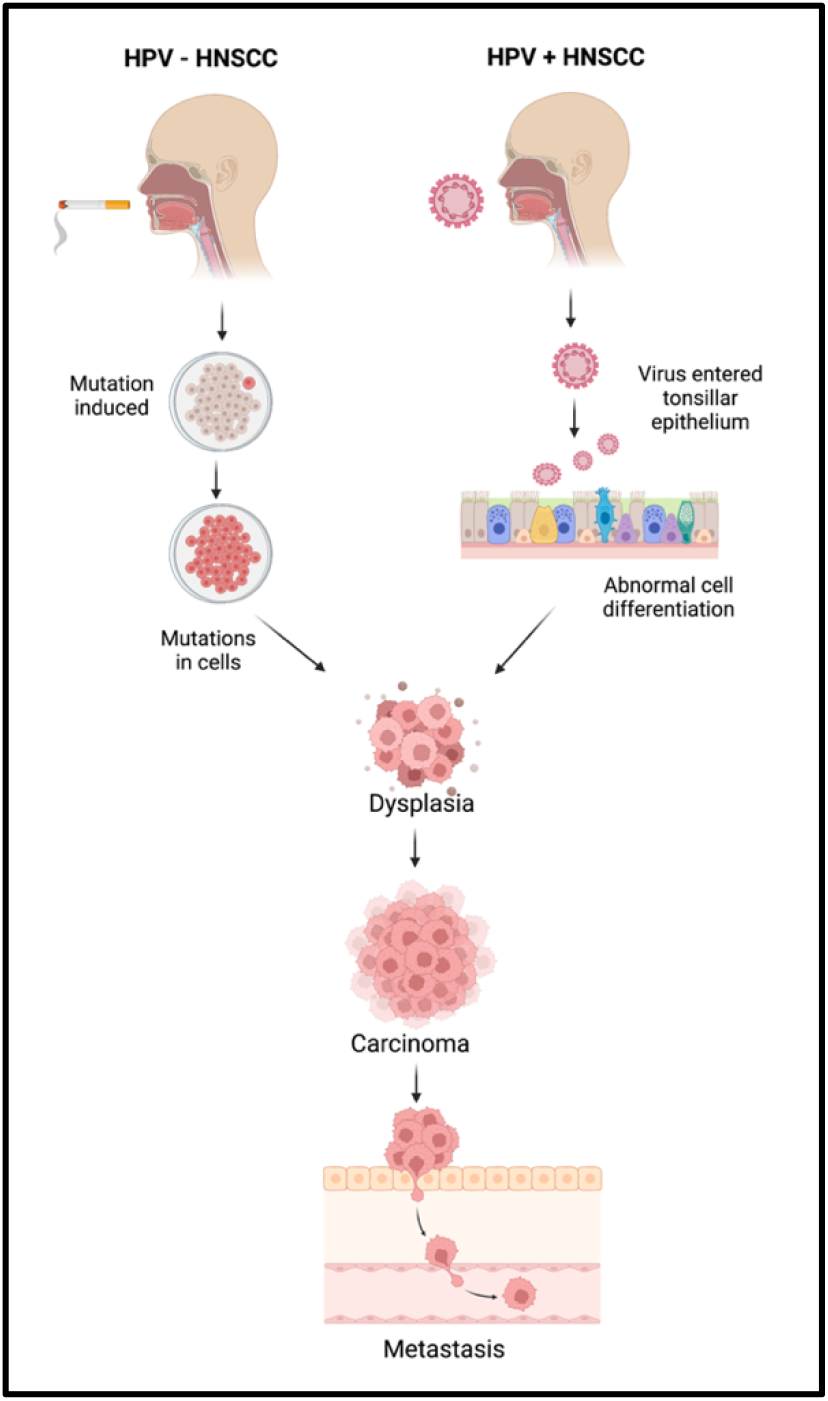
Shows mechanisms of head and neck squamous cell carcinoma (HNSCC) for HPV-positive and HPV-negative HNSCC patients.

Even after meticulous and selective therapy, the odds of survival are decreased by the fact that most cases of head and neck cancer are detected at advanced (late) stages. The traditional diagnosis of HNSCC is based on the physical examination, radiological investigation, and histological analysis of the tissue sections obtained from biopsies or surgical resections. These procedures can take a lot of time and are susceptible to mistakes in observation or interpretation, which can lead to discrepancies in cancer grading and prognostication [4]. In addition to this, most of the HNSCC cancers are detected at a later stage. The reasons range from limited symptomatology in early-stage patients, swift progression from early to advanced stage, indistinctive diagnostic characteristics, and imprecise history information [5].

Identification of molecular biomarkers of HNSCC can lead to early diagnosis of this cancer and can also help in preventive management of HNSCC. The cancer biomarkers not only influence diagnosis but they also have the potential to improve the treatment outcomes using targeted therapy. The currently known biomarker of HNSCC is PD-L1 which is commonly used in treatment decision making in advanced stage of HNSCC. It has a moderate predictive value and has many limitations due to the lack of standardization and highly dynamic nature of PD-L1 expression. Currently, there are no any other FDA approved molecular biomarkers for HNSCC diagnosis or prognosis [5].

In this study, we made an attempt to identify genetic biomarkers for HNSCC using single-cell sequencing data. On the basis of the 100 genetic biomarkers identified in this study, we have developed a method that can predict the HNSCC cancer along with HPV+ or HPV-status. Single-cell data collected from individual cells using next generation sequencing methods provides a better knowledge of the activity of a single cell in relation to its microenvironment [6]. Cell-to-cell variation can be revealed by single-cell sequencing of RNA or epigenetic alterations, which may aid the populations in quickly adapting to new circumstances [7]. The significance of gene mosaicism, as well as intra-tumor genetic heterogeneity in the genesis of cancer or response to therapy, can be uncovered by single-cell precision [8]. Single-cell technology makes it possible to detect molecular alterations in individual cancer cells. This can increase the research of more specialized biomarkers with excellent resolution, leading to the development of a complete landscape of distinct cell types within tumors [9]. The full workflow of this study is described in Figure 2.

**Figure 2:**
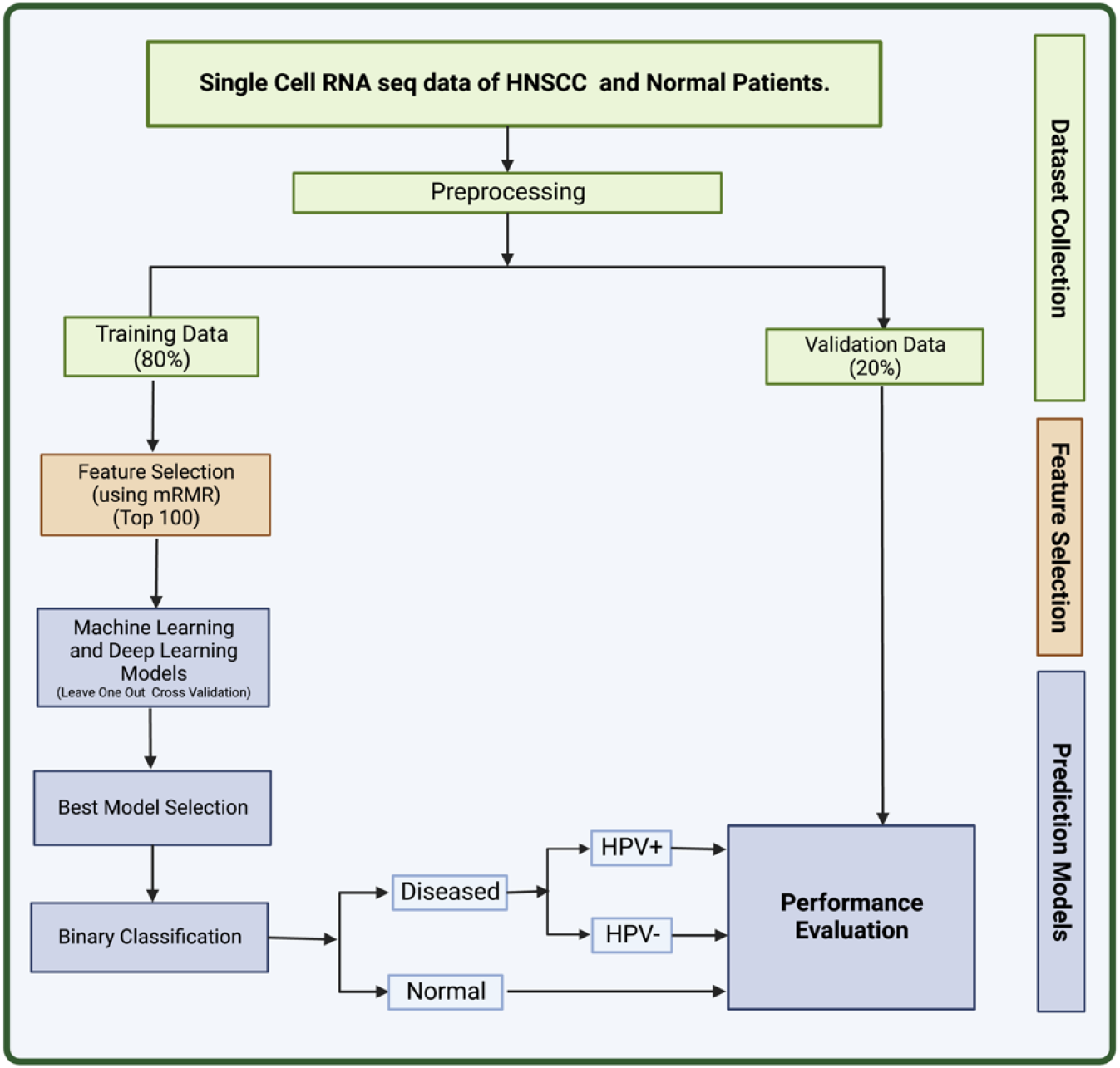
The full workflow of the study

## Material and Methods

### Data Collection

We retrieved the dataset used in this study (GSE181919) from National Centre for Biotechnology Information’s (NCBI) Gene Expression Omnibus (GEO) [10,11]. The dataset contains single-cell RNA sequencing data from 37 tissue specimen. The tissue specimen comprises 9 normal, 4 leukoplakia, 20 primary cancer (HNSCC), and 4 metastatic tumor (HNSCC) tissues. This dataset used Illumina HiSeq 400 as the platform for scRNA sequencing. The data processing methods include alignment and generation of barcode matrices using CellRanger. The information on whether the samples were derived from HPV+ or HPV-patients was derived from the metadata provided on GEO. The 80% of this dataset was used to train machine learning (ML) and deep learning (DL) models and 20% was used as validation set.

### Data Pre-processing

After the retrieval of data from GEO, we processed the data using in-house python scripts. Firstly, we converted the sparse data into a matrix and removed insignificant columns from our training data. The genes that had no mapped readings to more than 80% of the cells were eliminated, and cells containing zeroes were filtered leading to 2604 genes. The sequencing depth affects the range of values for the features, which necessitates normalizing the count data before doing any sort of analysis. Hence, we performed counts per million (CPM) normalization and log transformation on the data using scanpy package in python [12].

### Feature Selection

We applied feature selection to the set of 2604 genes obtained after pre-processing to obtain a set of genetic biomarkers for HNSCC. This was achieved using mRMR (Minimum Redundancy and Maximum Relevance) feature selection algorithm. mRMR selects a subset of features that have the least correlation amongst themselves but high correlation with the output class. The advantage of using this method is that it provides with a small set of features with high predictive potential. The redundancy between genes is taken into account in this technique in addition to the relationship between samples and genes. The most relevant feature will be considered out of the numerous identical features. We used the value K = 100 for mRMR to extract 100 most relevant genes for the prediction of HNSCC [13]. This strategy has been previously demonstrated to be useful and often utilized in single-cell RNA sequencing analysis [14,15].

### Machine Learning Models

We have developed various machine learning (ML) models to classify between normal subjects and HNSCC patients. In addition, we have also classified HNSCC patients into HPV positive and HPV negative. These machine learning models include Extreme Gradient Boosting (XGB), Decision Tree (DT), K-Nearest Neighbors (KNN), Extra Trees (ET), Logistic Regression (LR), and Random Forest (RF) algorithms. Hyperparameter tuning was also used to optimise the parameters of these algorithms. The DT classifier is a supervised machine learning model that classifies the output by learning decision rules from input, the KNN classifier predicts on the basis of the maximum number of votes cast in support of the class that is closest to the nearest neighbouring data point, LR classifier uses a logistic function to calculate the likelihood of an event, XGB Classifier is a distributed gradient-boosted decision tree machine learning package that offers simultaneous tree boosting, and RF classifier trains a number of decision trees to produce a single tree. A technique for ensemble supervised machine learning that makes use of decision trees is called extra trees.[16–21]. These methods have previously been used in many studies [22–25].

### Deep Learning Models

Along with the ML models, we have also applied deep learning classification technique – Artificial Neural Network (ANN) to classify the data [26]. In this method, networks are composed of multiple layers, and each layer has a number of nodes (or neurons) that support decision making. The model architecture of ANN used in this study includes three hidden layers and an output layer. A dropout of 0.5 is implemented at each step to lessen the overfitting by neural network. Biological neuron networks served as the basis for this strategy. Artificial neurons, which are constructed from a network of connected units or nodes and are conceptually similar to the neurons in the human brain, are used to build ANNs. They consist of several layers, and inside each layer there are multiple nodes (or neurons) that support decision-making. The anticipated label (Diseased or Normal) of the sample is the final result. The final result classifies the samples into HNSCC positive or negative, and if found HNSCC positive then whether the patient is HPV positive or negative is identified.

### Cross Validation

The dataset was primarily composed of training data, which made up 80% of it and validation set, which made up the remaining 20%. In the LOOCV (Leave One out Cross Validation) approach, the whole training set is separated into N equivalent folds using the LOOCV technique, with (N-1) being utilized for training and the single fold being used for testing. Each fold serves as testing data for the technique’s N iterations. The overall performance was calculated as the mean of N iterations. This is a common practice in many types of studies [27,28].

### Evaluation Parameters

To evaluate the efficacy of various prediction models, we employed a number of evaluation indicators. In this study, we used both threshold-independent and threshold-dependent parameters.

To calculate threshold-dependent characteristics like sensitivity (Sens), specificity (Spec), precision, F1-Score, and accuracy (Acc), we utilised the following formulae. We also used the conventional threshold-independent parameter Area Under the Curve (AUC) to assess the performance of the models.

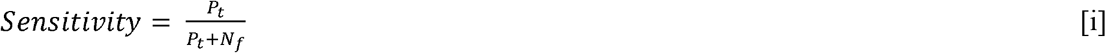

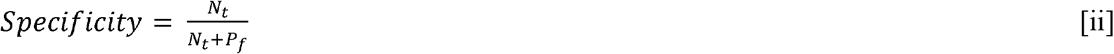

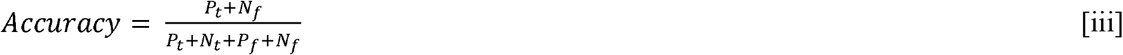

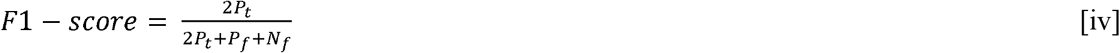

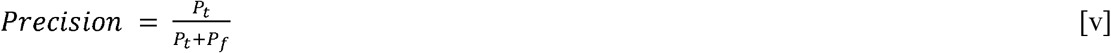

Where, *P*_*t*_ is true positive, *N*_*t*_ is true negative, *P*_*f*_ is false positive, and *N*_*f*_ is false negative.

## Results

### Feature Selection

We applied a feature selection technique called mRMR to obtain a list of highly relevant features (genes) for the detection of HNSCC samples from a set of 2604 genes that were obtained after data pre-processing [29]. We obtained a subset of 100 genes that were able to classify HNSCC and non-HNSCC samples as well as HPV+ and HPV-samples correctly. The list of selected 100 genes along with their p-values is given in Supplementary Table S1.

### Model Performance for HNSCC vs non-HNSCC

We applied various ML models like DT, RF, ET, XGB, and KNN on our dataset, where we used 80% of dataset GSE181919 for training, 20% of dataset GSE181919 as validation set, and. It was observed that machine learning models were able to achieve an AUROC of 0.85 (XGB, ET) on the validation set. In order to increase the AUROC, we applied DL algorithm – ANN on the dataset and observed that the AUROCs increased to 0.91 for the validation set. The complete results for the ML and DL performances are given in Table 1.

**Table 1:** Performance of ML and DL models for the classification of HNSCC patients and normal subjects.

| Training Set |  |  |  |  |  |  |  |
| --- | --- | --- | --- | --- | --- | --- | --- |
| Models | Accuracy | MCC | AUROC | Sensitivity | Specificity | Precision | F1 Score |
| Decision Tree | 0.93 | 0.85 | 0.93 | 0.95 | 0.91 | 0.94 | 0.94 |
| Random Forest | 0.96 | 0.92 | 0.96 | 0.98 | 0.94 | 0.96 | 0.97 |
| Logistic Regression | 0.92 | 0.84 | 0.92 | 0.95 | 0.88 | 0.92 | 0.94 |
| XGB | 0.92 | 0.83 | 0.92 | 0.92 | 0.92 | 0.95 | 0.93 |
| ExtraTree | 0.97 | 0.80 | 0.90 | 0.91 | 0.89 | 0.91 | 0.91 |
| KNN | 0.93 | 0.85 | 0.92 | 0.96 | 0.89 | 0.93 | 0.94 |
| Deep Learning Model | 0.99 | 0.93 | 0.97 | 0.98 | 0.96 | 0.97 | 0.98 |
| Validation Set |  |  |  |  |  |  |  |
| Models | Accuracy | MCC | AUROC | Sensitivity | Specificity | Precision | F1 Score |
| Decision Tree | 0.85 | 0.70 | 0.83 | 0.97 | 0.70 | 0.80 | 0.88 |
| Random Forest | 0.85 | 0.71 | 0.83 | 0.99 | 0.68 | 0.79 | 0.88 |
| Logistic Regression | 0.79 | 0.60 | 0.77 | 0.98 | 0.56 | 0.73 | 0.84 |
| XGB | 0.86 | 0.73 | 0.85 | 0.98 | 0.71 | 0.81 | 0.89 |
| ExtraTree | 0.86 | 0.74 | 0.85 | 0.99 | 0.71 | 0.81 | 0.89 |
| KNN | 0.83 | 0.68 | 0.81 | 0.98 | 0.65 | 0.77 | 0.87 |
| Deep Learning Model | 0.92 | 0.82 | 0.91 | 0.94 | 0.89 | 0.94 | 0.94 |

### Model Performance for HPV+ vs HPV-

After classification of samples as HNSCC or non-HNSCC, we attempted to classify whether an HNSCC sample belonged to an HPV+ or HPV-class. Hence, we developed ML and DL models to further classify the HNSCC samples as HPV+ and HPV-. The maximum AUROC achieved by ML models was 0.81 (XGB) for the validation set. After employing ANN classifier to the data, it was observed that the AUROC increased to 0.84 for the validation set. The results for HPV+ and HPV-classification from HNSCC patients are summarized in Table 2.

**Table 2:** Performance of ML and DL models for the classification of HPV+ and HPV-patients from HNSCC patients.

| Training Set |  |  |  |  |  |  |  |
| --- | --- | --- | --- | --- | --- | --- | --- |
| Models | Accuracy | MCC | AUROC | F1 Score | Sensitivity | Specificity | Precision |
| Decision Tree | 0.75 | 0.50 | 0.75 | 0.78 | 0.79 | 0.71 | 0.77 |
| Random Forest | 0.82 | 0.63 | 0.81 | 0.84 | 0.87 | 0.75 | 0.81 |
| Logistic Regression | 0.84 | 0.67 | 0.84 | 0.86 | 0.86 | 0.82 | 0.85 |
| XGB | 0.77 | 0.52 | 0.76 | 0.79 | 0.81 | 0.71 | 0.77 |
| ExtraTree | 0.84 | 0.68 | 0.84 | 0.86 | 0.88 | 0.8 | 0.84 |
| KNN | 0.80 | 0.59 | 0.79 | 0.82 | 0.84 | 0.75 | 0.8 |
| Deep Learning Model | 0.991 | 0.980 | 0.995 | 0.992 | 0.989 | 0.993 | 0.995 |
| Validation Set |  |  |  |  |  |  |  |
| Models | Accuracy | MCC | AUROC | F1 Score | Sensitivity | Specificity | Precision |
| Decision Tree | 0.69 | 0.35 | 0.65 | 0.74 | 0.76 | 0.59 | 0.72 |
| Random Forest | 0.84 | 0.88 | 0.80 | 0.83 | 0.91 | 0.92 | 0.94 |
| Logistic Regression | 0.80 | 0.54 | 0.75 | 0.85 | 0.9 | 0.61 | 0.81 |
| XGBClassifier | 0.82 | 0.61 | 0.81 | 0.86 | 0.87 | 0.74 | 0.86 |
| ExtraTree Classifier | 0.46 | -0.11 | 0.76 | 0.52 | 0.58 | 0.33 | 0.47 |
| K Neighbours classifier | 0.49 | -0.06 | 0.55 | 0.55 | 0.58 | 0.38 | 0.52 |
| Deep Learning Model | 0.84 | 0.70 | 0.83 | 0.88 | 0.98 | 0.68 | 0.79 |

### Gene ontology

The Gene Ontology (GO) encapsulates our understanding of the biological world in three ways: molecular function, cellular component, and biological process [30,31]. 100 genes that may serve as potential biomarkers of HNSCC were retrieved once mRMR analysis was complete. On these 100 retrieved genes, we next ran Gene Ontology (GO) enrichment analysis using PantherDB to map the biological functions of the chosen genes [32]. The findings of the GO enrichment analysis are displayed in Table 4. We see that the majority of genes have a role in the catalytic and binding activities of many metabolic processes. ATP-dependent activity, molecular function regulator, molecular transducer, structural molecule activity, translation regulator activity, transcription regulator, and transporter activity are additional activities linked to the reported genes. The genes and their roles are described in Table 3.

**Table 3:** Gene Ontology (GO) enrichment results for top 100 selected genes.

| GO Term | Activity | Genes |
| --- | --- | --- |
| (GO:0140657) | ATP-dependent activity | ABCA8 |
| (GO:0005488) | binding | UBC, ADAR, ICAM3, TXNIP, FXYD1, TIGIT, EEF1A1, RPLP1, LIMD2, PSME1, RPS26, EZR, HLA-C, RHOG, FIGF, LTBP4, OAZ1, B2M, RAC2, TIMP3, FHL1, UBB, HLA-B, BMP4, ARPC3, RELB, EBF1, RHOH, ADH1B, RPS19, IL2RG, GSN, SOD3, GDF10, SPARCL1, NOVA1, CREM, REL, HLA-A |
| (GO:0003824) | catalytic activity | CYBRD1, DUSP4, ADAR, PIM3, EEF1A1, RPLP1, PSME1, ISG20, ATP5E, RHOG, OAZ1, RAC2, TIMP3, PTGIS, ABCA8, RHOH, ADH1B, SOD3, PLPP3 |
| (GO:0098772) | molecular function regulator | FXYD1, RPLP1, PSME1, FIGF, OAZ1, TIMP3, BMP4, GDF10 |
| (GO:0060089) | molecular transducer activity | FIGF, BMP4, IL2RG, GDF10 |
| (GO:0005198) | structural molecule activity | RPS15A, RPLP1, RPS26, RPS11, RPS15, RPS19 |
| (GO:0140110) | transcription regulator activity | RELB, EBF1, CREM, REL |
| (GO:0045182) | translation regulator activity | EEF1A1 |
| (GO:0005215) | transporter activity | FXYD1, ATP5E, AQP7, SLC16A3, ABCA8 |

## Discussion

One of the heterogeneous diseases, HNSCC affects the head and neck region, namely the oral cavity, paranasal sinuses, larynx, nasal cavity, hypopharynx, and oropharynx. It is described by malignant and uncontrollable cell proliferation in these locations [33]. Advancement in the sequence technology allows the researchers to find various biomarkers such as diagnostic, predictive, and prognostic biomarkers. These biomarkers helps in better understanding of the disease as well as may aids in designing novel and effective diagnosis and treatment. A biomarker is described as a biological molecule present in the blood, other body fluids, as well as in tissues, that serves as a sign of a normal or aberrant process, a condition, or a disease by the National Cancer Institute (NCI). To determine how effectively the body will react to an illness or condition medication, a biomarker could well be utilized [33]. This study aims to find out a set of potential biomarkers from single-cell genomic data of head and neck cancer patients that can classify HNSCC patients and normal individuals with reliable accuracy. In addition, we have also attempted to classify HNSCC patients as HPV+ or HPV-. The biomarkers identified in this study could aid in the early diagnosis and screening of HNSCC.

To categorize NC and HNSCC disease cells from their single-cell RNA seq data, we employed a variety of machine learning models, including an ANN deep learning model. We also further tried to categorize the diseased patients into HPV+ and HPV-. We trained the model with 80% of the dataset GSE181919, 20% of the dataset GSE191919 as validation set. The datasets were originally quite extensive and had a significant number of features. During the preprocessing step, the feature count was decreased to a shallow level of 2604 genes (features). One of the feature selection techniques known as mRMR was used to obtain the limited set of features which could be helpful in categorizing the samples because many characteristics were correlated. The top 100 genes with the least amount of redundancy and the most relevance were extracted from these 2604 genes (features) using mRMR. Furthermore, 100 genes (features) separated the HNSCC patients from NC with an accuracy of around 92%, an AUROC of 0.91 in the validation set. Whereas in the case of HPV classification, the metrics obtained were, AUROC 0.83 and 98% accuracy on the validation set. For the detection and categorization of biomarkers, ANN has proven to be an effective technique among all machine learning models.

After obtaining the top 100 most relevant genes for the classification of HNSCC, we performed Gene Ontology (GO) enrichment analysis using PantherDB and most of the genes were observed to be related to catalytic and binding activities [32]. Some of them also had a role in other essential processes like ATP-dependent activity, molecular function regulator, molecular transducer, structural molecule activity, translation regulator activity, transcription regulator, and transporter activity. Many of the genes identified in this study have been previously linked to HNSCC in earlier studies. The gene PLAC9’s overexpression has been reported in connection with the inhibition of cell growth regulation and has also been reported in connection with cancers such as ovarian cancer and breast cancers as prognostic biomarkers [34]. Gene “ACKR1”, along with other 3 genes in a study, was reported to be downregulated in HNSCC patients, which was correlated with poor prognosis (p<0.05) [35]. Also, gene “AQP7”, which is involved in physiologically functional cell migration, was upregulated in MSR of patients with ten tumors [36]. Whereas, gene FXYD1 was reported to be downregulated in the cancer samples, while FXYD4 and FXYD5 were overexpressed (P<0.05, fold change>1.5) [37]. In a study on cancer cells, it was observed that BTG1 gene overexpression was linked to tumor growth or lung metastasis, inhibited proliferation, and induced differentiation in different types of cancer cells [38]. Also, mutations occurring in different genes, including B2M, CDKN2A, is found to be related with the occurrence and development of tumors in Head and neck cancer patients [39]. Genes such as MFAP4, CD37, CXCL12, ADH1B, SOD3, SCARA5, ANGPTL1, FHL1, F10, CXCR4, MEG3, TXNIP, GDF10, and ABI3BP are shown to be downregulated because they operate as potential tumor suppressor genes, inhibiting tumor cell proliferation, invasion, and migration while also promoting apoptosis [40]. By controlling the expression of miR-421 and E-cadherin, MEG3 long-encoding RNA inhibits the development of head and neck squamous cell carcinoma. However, additional research into MEG3’s downstream mechanism in controlling the molecular process of epithelial-mesenchymal transformation (EMT) in head and neck squamous cell carcinoma (HNSCC) development is required [41]. Growth differentiation factor-10 (GDF10), also known as BMP3b, is a tumor suppressor that belongs to the transforming growth factor-b (TGF-b) superfamily [42]. CIB1, PIM3, SLC16A3, VOPP1, BMP4, TIGIT, ADAR, and LRRN4CL are studied as upregulated genes [2,43–51]. A complex that is important in the keratinocyte-intrinsic immune response to human papillomaviruses (-HPVs) is formed when CIB1 interacts with the EVER1, and EVER2 proteins [43,44]. It has been observed that nearly all primary HNSCCs express at least one PIM kinase member at high levels [2]. Immunological checkpoint T cell immunoreceptor with immunoglobulin and ITIM domain (TIGIT) is essential for immune suppression. However, it has a connection to genetics and epigenetics, and a role in tumor immunity [48]. The transforming growth factor (TGF) superfamily includes extracellular signaling molecules known as bone morphogenetic proteins (BMPs), which are known to control cell proliferation, differentiation, and motility, particularly during development. Functional research shows that, particularly in HNSCC cancer, has connected BMP4 to the encouragement of cell migration and the suppression of cell proliferation [47].

Overall, most of the genes which were obtained from our study have been reported as promising candidate for biomarkers in various studies [2,35–40]. However, some genes have not yet been reported in connection with Head and Neck cancer. These genes may require further investigation and study. These genes may act as novel findings which could help in diagnose patients with Head and neck cancer. In order to help the scientific community, we created a Python package called “HNSCPred” based on the aforementioned work (https://webs.iiitd.edu.in/raghava/hnscpred/). To fully understand how the discovered genes impact and contribute to the progression of HNSCC disease, further clinical investigations on these genes are necessary.

## Author’s contributions

AJ, AD, SP, and AS collected and processed the data. AJ, AD, SP, and AS implemented the algorithms, developed the prediction models and python package. AA and GPSR prepared the manuscript. AA developed the web interface for the tool. GPSR conceived and coordinated the project. All authors have read and approved the final manuscript.

## Acknowledgements

The authors are thankful to the Council of Scientific and Industrial Research (CSIR), Department of Biotechnology (DBT), and Department of Science and Technology (DST) for providing fellowships and financial support. The authors are also thankful to the Department of Computational Biology, IIITD, New Delhi for its infrastructure and facilities. We thank the Department of Biotechnology (DBT) for providing an infrastructure grant to the institute (Grant BT/PR40158/BTIS/137/24/2021). We would like to acknowledge that figures were created using BioRender.

## Data Availability Statement

All the codes and dataset generated in this study are available at https://webs.iiitd.edu.in/raghava/hnscpred/

